# Gauss-power mixing distributions comprehensively describe stochastic variations in RNA-seq data

**DOI:** 10.1101/194118

**Authors:** Akinori Awazu, Takahiro Tanabe, Mari Kamitani, Ayumi Tezuka, Atsushi J. Nagano

## Abstract

**Motivation:** Gene expression levels exhibit stochastic variations among genetically identical organisms under the same environmental conditions. In many recent transcriptome analyses based on RNA sequencing (RNA-seq), variations in gene expression levels among replicates were assumed to follow a negative binomial distribution although the physiological basis of this assumption remain unclear.

**Results:** In this study, RNA-seq data were obtained from *Arabidopsis thaliana* under eight conditions (21–27 replicates), and the characteristics of gene-dependent distribution profiles of gene expression levels were analyzed. For *A. thaliana and Saccharomyces cerevisiae,* the distribution profiles could be described by a Gauss-power mixing distribution derived from a simple model of a stochastic transcriptional network containing a feedback loop. The distribution profiles of gene expression levels were roughly classified as Gaussian, power law-like containing a long tail, and mixed. The fitting function predicted that gene expression levels with long-tailed distributions would be strongly influenced by feedback regulation. Thus, the features of gene expression levels are correlated with their functions, with the levels of essential genes tending to follow a Gaussian distribution and those of genes encoding nucleic acid-binding proteins and transcription factors exhibiting long-tailed distributions.

**Availability:** Fastq files of RNA-seq experiments were deposited into the DNA Data Bank of Japan Sequence Read Archive as accession no. DRA005887. Quantified expression data are available in supplementary information.

**Supplementary information:** Supplementary data are available at *Bioinformatics* online.

## 1 Introduction

Stochastic variations in gene expression—known as gene expression noise or phenotype fluctuation—have been observed among individuals in a genetically identical population under the same environmental conditions (Elowitz et al., 2002; Furusawa et al., 2005; Golding et al., 2005; Kaern et al., 2005; Newman et al., 2006; Chang et al., 2008; Konishi et al., 2008; Taniguchi et al., 2010; So et al., 2011; Silander et al., 2012; Woods, 2014). Such variations are thought to be important for maintaining the pluripotency of embryonic stem cells, cell fate decisions, and cellular differentiation in multicellular organisms (Mitsui et al., 2003; Kaneko, 2006; Kalmar et al., 2009; Ochiai et al., 2014). In rice, genes related to stress responses exhibited larger variations than those involved in other processes (Nagano et al., 2012). Furthermore, recent studies in *Escherichia coli*, the budding yeast *Saccharomyces cerevisiae*, and *Arabidopsis thaliana* have reported that the magnitude of gene expression noise is positively correlated with plasticity—i.e., the variation in expression levels due to mutation or environmental change (Sato et al., 2003; Blake et al., 2003; Landry et al., 2007; Choi and Kim, 2008; Choi and Kim, 2009; Tirosh and Barkai, 2008; Lehner, 2010; Lehner and Kaneko, 2011; Bajic and Poyatos, 2012; Singh, 2013, Hirao et al., 2015).

Recent gene expression analyses with sufficiently large replicates have shown that in organisms as diverse as *E. coli* and mammals, fluctuations in protein expression level for a given gene follow a log-normal distribution (Sato et al., 2003; Furusawa et al., 2005; Chang et al., 2008; Konishi et al., 2008). The closely related Frechet distribution was also proposed to describe variations in gene expression levels in *E. coli* and *S. cerevisiae* (Salmann et al., 2012). On the other hand, mathematical modeling of protein expression in *E. coli* suggested that such variations were more closely approximated by a gamma distribution, which is often considered as log-normal (Friedman et al., 2006; Taniguchi et al., 2010).

In many recent high-throughput RNA sequencing (RNA-seq) studies (Mortazavi et al., 2008; Nagalakshmi et al., 2010), variations in gene expression (transcription) levels among replicates were assumed to follow a negative binomial (NB) distribution (Marioni et al., 2008; Robinson et al, 2008; Rapaport et al., 2013; Gierlin’ski et al., 2015; Schurch et al., 2016). An analysis of RNA-seq data from a two-condition, 48-replicate experiment using *S. cerevisiae* revealed that variations in expression levels for each gene conformed to both log-normal and NB distributions (Gierlin’ski et al., 2015). Beta-binomial and Benford distributions have been proposed for fitting gene expression data obtained by RNA-seq (Smith et al., 2016; Karthik et al., 2016). However, the physiological basis of these distributions and the significance of associated parameters remain unclear. Furthermore, it is not known whether such model distributions are applicable to any genes in any organism, especially multicellular organisms.

Gene expression noise in plants has been investigated in rice and *Arabidopsis* (Nagano et al., 2012; Shen et al., 2012, Hirao et al., 2015). However, recent studies were based on transcriptome data from experiments with few replicates (Maruyama-Nakashita et al., 2005; Nemhauser et al., 2006; Kilian et al., 2007; Goda et al., 2008; Less and Galili, 2008; Wittenberg et al., 2012), which limited the inferences that could be made regarding the distribution characteristics of gene expression levels. In the present study, we analyzed RNA-seq data for *A. thaliana* under eight conditions (21–26 replicates) to determine distribution profiles of gene expression noise (phenotype fluctuation) among individuals in a homogeneous population. We fitted the distribution profiles with a novel distribution function that we termed the Gauss-power (G-P) mixing distribution, which was derived from a simple stochastic transcriptional network model containing a feedback loop. The expression of genes showing a long-tail distribution was strongly influenced by a feedback mechanism; moreover, variations in gene expression levels were correlated with average expression levels and gene function.

## 2 Results

### 2.1 Analysis of Arabidopsis RNA-seq data

RNA-seq data from 7- and 22-day-old *Arabidopsis* shoots cultured under a 12:12-h light/dark cycle were obtained 1, 7, 13, and 19 h after the lights were turned on. There were 21 to 27 replicates for each condition. In total, 189 individual plants were analyzed by RNA-seq. We obtained 8.4 million reads on average; 1 sample with fewer than 1 million reads mapped to genes was omitted from subsequent analyses. The expression level of each gene was defined as the number of reads mapped to each gene per 1 million reads (Table S1). For each condition, we examined the distribution profiles of expression levels of ∼10,000 genes (Table 1) whose expression levels could be regarded as stationary (see Materials and Methods).

**Table 1.** Number of replicates in the RNA-seq experiment and number of analyzed genes for each condition of *Arabidopsis*

| Condition | Age (days) | Time after light (h) | Replicate | No. of analyzed genes |
| --- | --- | --- | --- | --- |
| 7-1 | 7 | 1 | 21 | 10,499 |
| 7-7 | 7 | 7 | 21 | 9287 |
| 7-13 | 7 | 13 | 25 | 10,760 |
| 7-19 | 7 | 19 | 24 | 10,735 |
| 22-1 | 22 | 1 | 22 | 9619 |
| 22-7 | 22 | 7 | 24 | 12,109 |
| 22-13 | 22 | 13 | 24 | 12,810 |
| 22-19 | 22 | 19 | 27 | 11,338 |

### 2.2 Cluster analysis of rank-expression level distribution (RED) profiles of genes

RED profiles were obtained for each gene under each condition (harvest time and age of plant) (Fig. 1). For most genes, the REDs showed typical profiles but were very noisy. However, it is expected that groups of genes whose expression is similarly regulated would have similar RED profiles. We performed a cluster analysis to extract the essential properties of the RED profiles for each gene (see Materials and Methods).

**Figure 1.**
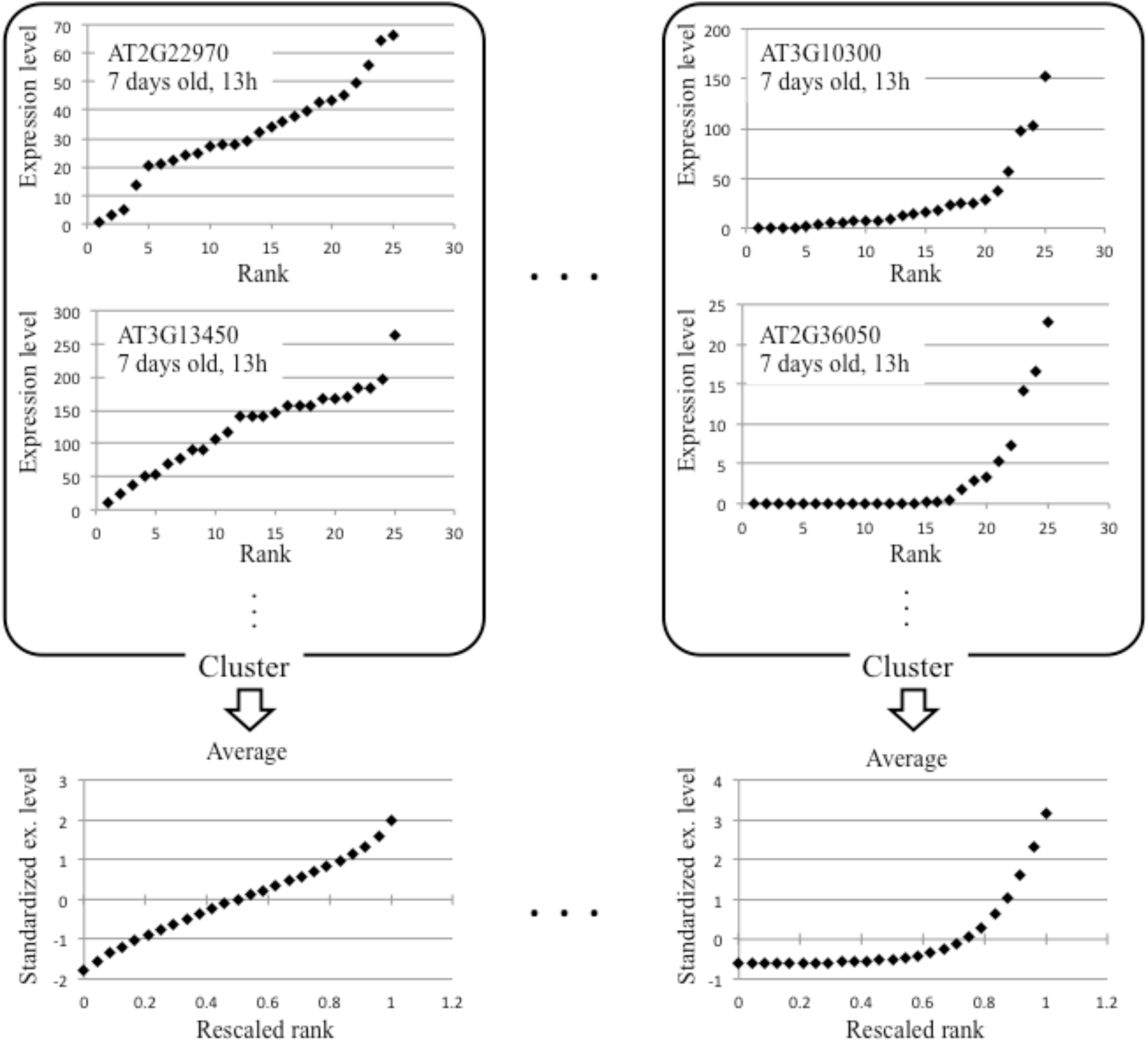
RED profiles of genes and average normalized RED profiles of their respective clusters. (Upper) RED profiles of indicated genes under specific conditions. As examples, results obtained from RNA-seq data at 13 h for 7-day-old *Arabidopsis* are shown. Similar RED profiles were grouped by cluster analysis. (Lower) Average normalized RED profiles for genes belonging to specific clusters.

For each condition, 12–15 clusters were obtained from normalized RED profiles (Tables S2 and S3), representing the relationship between standardized expression levels for mean = 0 and standard deviation = 1 and rescaled rank from 0 to 1. The average values of standardized expression levels of genes belonging to the same cluster were expected to reflect the essential features of their RED profiles (Fig. 1).

### 2.3 Inferences on probability density distribution profiles of gene expression levels

Since gene expression levels are non-negative, we analyzed normalized RED functions that were shifted such that the minimum value on the vertical axis was assumed to be 0 (referred to as shifted normalized RED function) instead of the previous normalized RED functions. The value of the vertical axis of this function represented the rescaled expression level. The inverse function of the shifted normalized RED function provided the probability distribution of rescaled expression levels (Fig. 2), whose derivative yielded the profiles of probability density distribution function of rescaled expression levels (PDL) for each cluster (Figs. 2 and S1–S8). The derivative of this function was estimated by differential approximation (see supplementary information S1). PDL profiles obtained from the clusters showed variable shape, including Gaussian and power law-like distributions.

**Figure 2.**
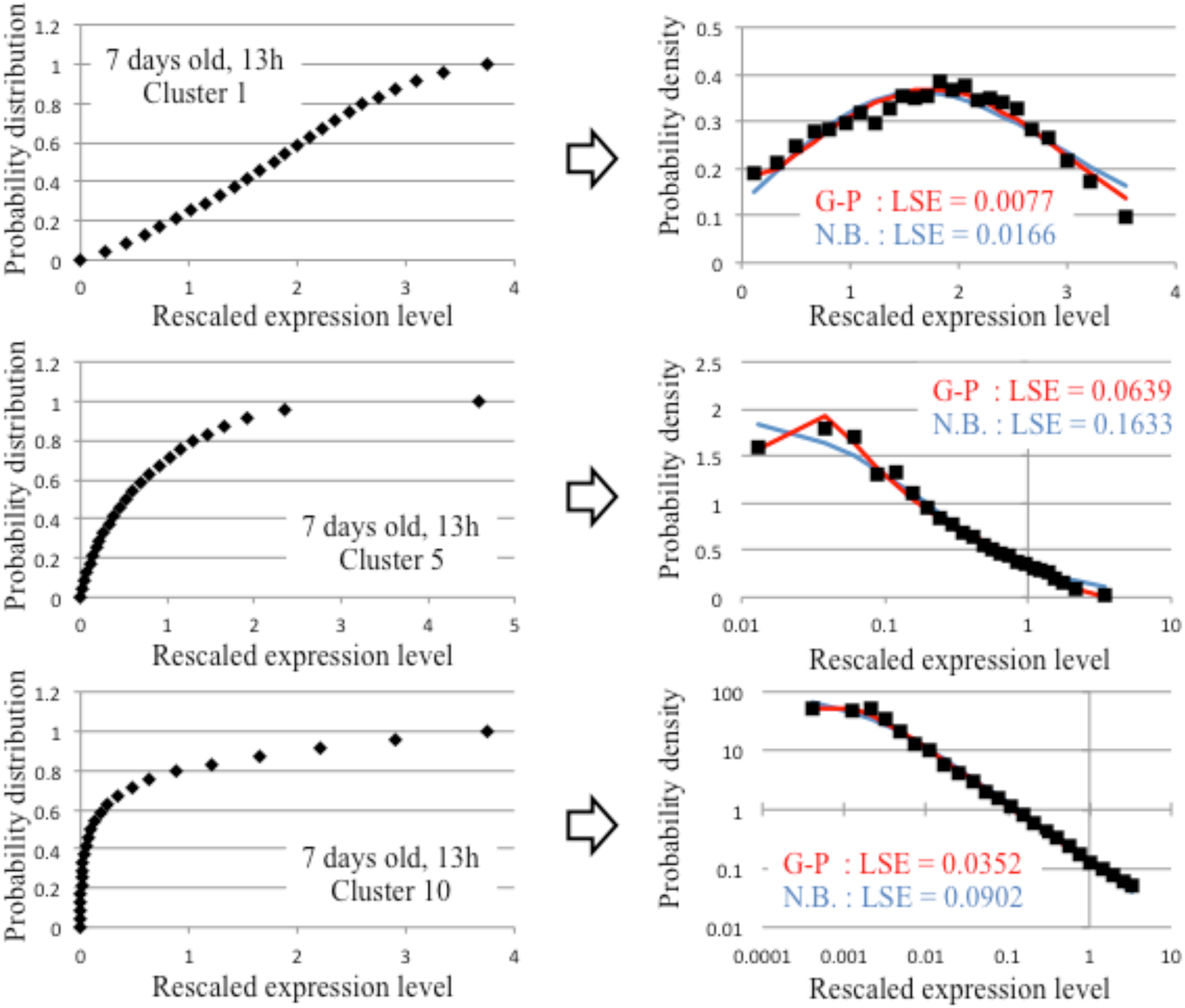
Examples of probability distributions and PDL profiles for indicated clusters. Examples of profiles of probability distribution of rescaled expression levels (left) and of PDL profiles (right) for three clusters. Red and blue represent curves fitted with the G-P and NB distribution functions, respectively. Least square error was estimated for the fitting curves.

### 2.4 G-P mixing distribution

The distribution profiles of gene expression levels were systematically classified based on the following mathematical model. A novel distribution function, which we refer to as G-P distribution, is described by equation 1.

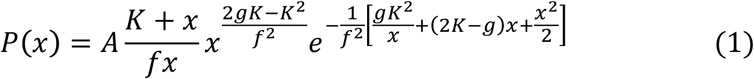

This equation is a fitting function of the distribution function of expression level *x* of the gene of interest *X*; the parameter *A* is a normalized coefficient; and *f*, *g*, and *K* are constants whose physiological significance is described below.

In general, gene expression levels are increased by activation and decreased by inhibition of upstream genes in a gene regulatory network. Furthermore, temporal changes in gene expression levels are directly or indirectly influenced by genes themselves, since the gene regulatory network includes many positive and negative feedback loops that are activated in a stochastic manner based on fluctuations in gene expression. Thus, a simplified model of the temporal change in the expression level *x* of gene *X* influenced by upstream genes and feedback regulation (Fig. 3) is given by equation 2:

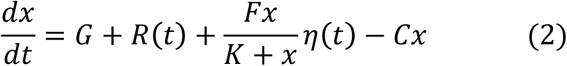

where *R(t)* and *η*(t) are assumed to be Gaussian white noise with 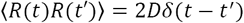 and 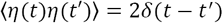; the parameters *G*, *K*, *F*, and *C* are ∼[average activation rate of *X* by upstream genes], ∼[average expression level of *X* required to induce maximum expression of downstream genes], ∼[magnitude of feedback effects], and [degradation rate of *X*], respectively; and P(x) is the steady-state probability distribution for simplified cases where D ? 0 where *g* = *G*/*C* and *f* = *F*/*C* (see supplementary information S2).

**Figure 3.**
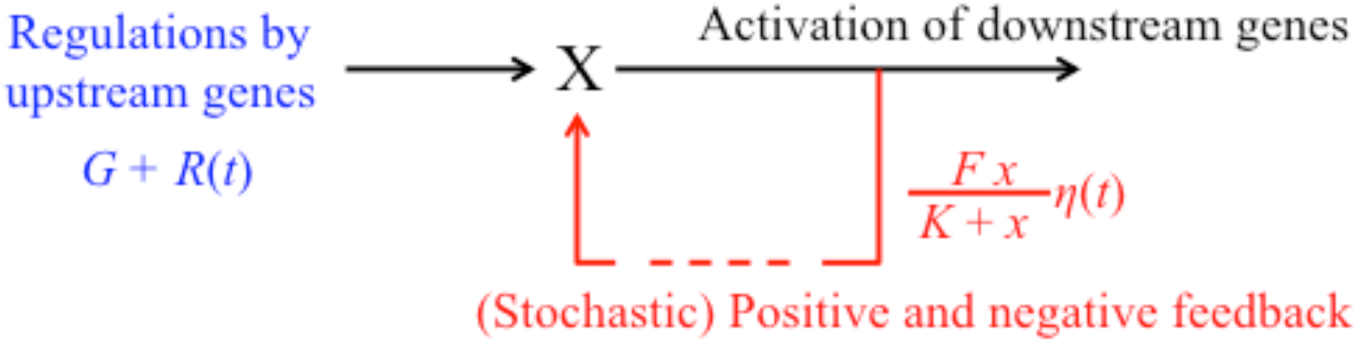
Illustration of a gene network model that fits a G-P distribution function. Gene *X* is regulated by upstream genes and by stochastic feedback.

### 2.5 Fitting of PDL profiles with the G-P mixing distribution

PDL profiles of each cluster under each condition were fitted with the G-P mixing distribution and (generalized) NB distribution according to equation 3:

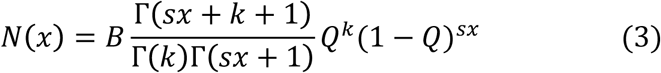

where Γ (*r)* is the gamma function and *B*, *s*, *k*, and *Q* are fitting parameters. Note that the parameter *s*—which is usually equal to 1—contributes to the generalization for various scales of x

The characteristics of PDL profiles for some clusters can be extracted from plots with a linear scale axis; however, it is more difficult to extract those of profiles with much larger maximum values and exhibit power law-like profiles, for which log-log plots seem more suitable when the maximum PDL value is greater than 3. In order to extract their detailed characteristics, PDL profiles were fitted using a typical least squares method for maximum PDL values < 3; PDL fitting parameters were chosen so as to minimize the sum of squared errors between log[PDL] and log[fitting functions] when the maximum PDL value was > 3. The results suggest that the G-P distribution has a least square error that is equal to or smaller than that of the NB distribution for PDL profiles of most clusters (Fig. 2 and Table S3). Therefore, in subsequent analyses the PDL profiles were classified according to a G-P distribution.

### 2.6 Classification of PDL profiles

When PDL profiles of each cluster were fitted to the G-P distribution function, some had *K* = 0, indicating that they were Gaussian, while others had *K* >> *g*, indicating that they were closer to a power law distribution. PDL profiles were classified as one of three types: Gaussian (*K* = 0), power law-like (*K* >> *g*), or mixed (*K ≈ g*) (Table S3). Here, *f* >> *g* as also obtained when *K* >> *g*, indicating that when the influence of feedback effects are large relative to the other mechanisms regulating gene expression, gene expression levels exhibit a long-tailed power law-like distribution.

Even for the same gene, PDL profiles varied depending on plant age and harvest time (Table S2). The ratio of occurrence of Gaussian, mixed, and power law-like distributions at four time points in younger plants (7 days old) was ∼ 3:3:4, while that of older plants (22 days old) was ∼45:29:26. High average expression levels were more frequently associated with a Gaussian as compared to a power law-like distribution; average expression levels and peak value of the frequency distribution were higher for the former than for the latter (Fig. 4).

**Figure 4.**
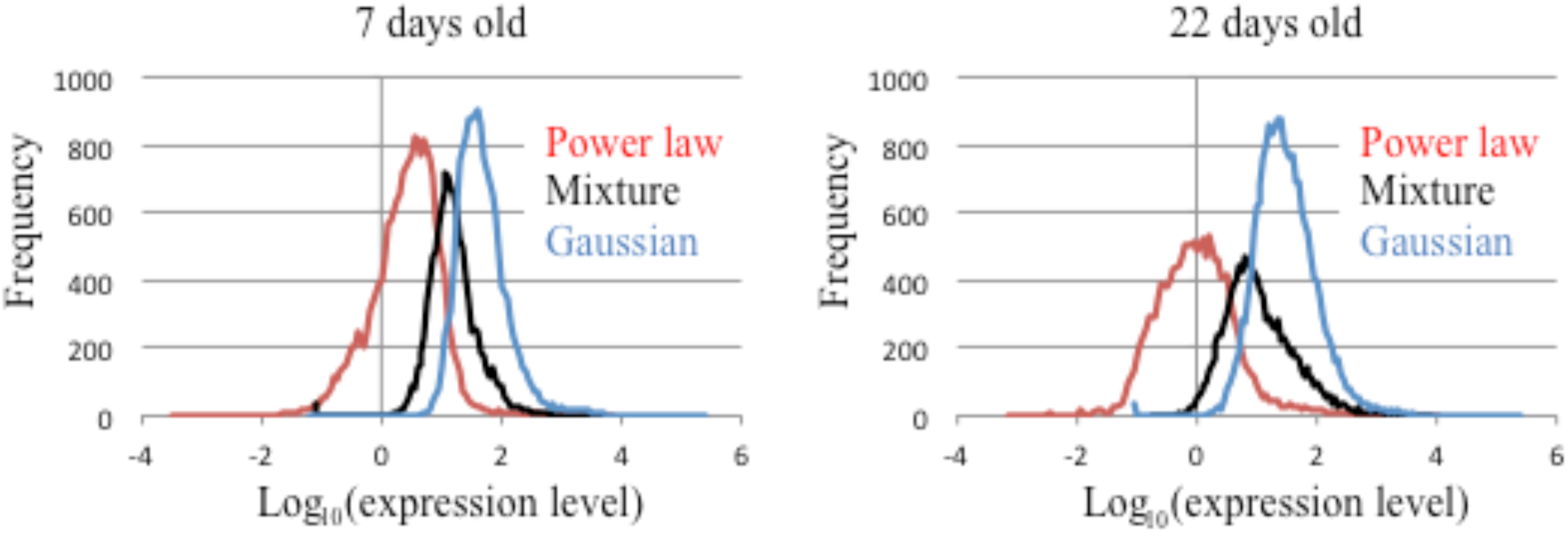
Frequency distributions of average log gene expression levels of gene groups exhibiting distinct PDL profiles. Frequency distributions of average log gene expression levels in cases of Gaussian (blue), mixed (black), and power law-like (red) distributions in 7-day-old (left) and 22-day-old (right) *Arabidopsis*. Differences in average log gene expression levels between Gaussian and mixed, and between mixed and power law-like distributions were significant (P < 0.01, t test) at both plant ages.

Gene function was also correlated with PDL profiles (Tables 2 and S4). For example, More than half of “essential genes” (Meinke et al., 2008)showed Gaussian PDL distribution at four time points of young plants and old plants. Furthermore, genes encoding important intracellular components and organelles and those associated with electron transport in metabolic pathways tended to show Gaussian PDL distributions. On the other hand, genes encoding transcription factors and nucleic acid-binding proteins mostly exhibited power law-like and mixed PDL distributions.

**Table 2.**
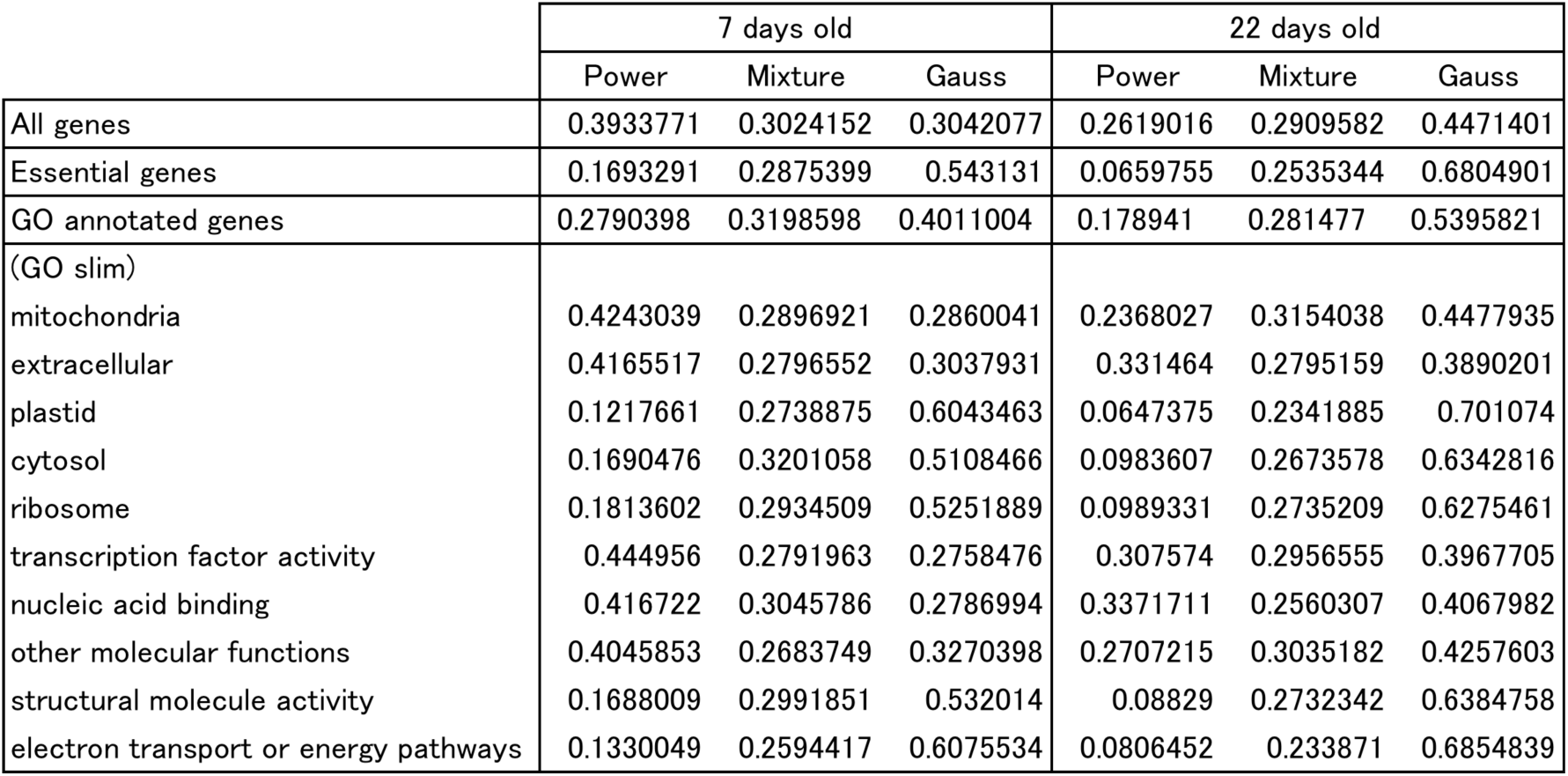
Relationships between gene groups classified according to function and ratio of occurrences of PDL profiles

## 3 Discussion

More than 20 replicates of *A. thaliana* gene expression data at four harvest times of 7- and 22-day-old shoots were obtained and the PDL profiles of each gene were analyzed. Most profiles could be fitted by the G-P distribution. There were three typical PDL profiles; namely, Gaussian, power law-like, and mixed.

The G-P distribution suggested that the various types of PDL profile were highly influenced by network topology, particularly a feedback loop regulating gene expression; for instance, gene groups showing a power law-like distribution were frequently influenced by a feedback mechanism, while this was rare for those exhibiting a Gaussian distribution. Furthermore, the PDL profiles of genes were correlated with their average expression levels and functions; gene groups classified as being essential for survival tended to exhibit Gaussian distributions, whereas those encoding transcription factors and nucleic acid-binding proteins mostly followed a non-Gaussian (i.e., power law-like or mixed) distribution. Furthermore, the expression levels of many genes classified as “unknown” exhibited power law-like distributions (Table S4), suggesting that their expression is predominantly modulated by feedback loops.

PDL profiles of gene expression levels were inferred from publicly available RNA-seq data derived from 48-replicate experiments of *S. cerevisiae* (Gierlin’ski et al., 2015) in the same manner as in the present study. The G-P as well as the NB distribution function fit the PDL profiles of *S. cerevisiae* (Fig. S9). However, long-tailed power law-like PDL profiles were not observed, unlike for *Arabidopsis* genes.

Even when the analysis was performed using 24-replicate data randomly selected from the 48-replicate dataset, the results were qualitatively similar to those described above, except that the number of clusters differed (Fig. S10). Although the number of replicates in the present study was smaller than that used in the earlier report, our results reflect the essential properties of the PDL profiles of *Arabidopsis* genes and are expected to apply to a larger number of replicates.

This study mainly focused on the steady-state probability distributions of gene expression levels. However, many *Arabidopsis* genes are regulated by circadian rhythm. Future studies must therefore address the extent to which the present model can be generalized to dynamic situations. Furthermore, intermittent temporal changes were observed in the expression of genes following a power law-like distribution, suggesting a transcriptional burst mechanism (Taniguchi et al., 2010; So et al., 2011; Munsky et al., 2012; Sanchez et al., 2013; Jones et al., 2014; Fujita et al., 2016) for genes whose expression is predominantly influenced by feedback. Such dynamic features of gene expression warrant more detailed analysis.

## 4 Materials and Methods

### 4.1 Plant growth conditions and RNA-seq

Seeds of *A. thaliana* (accession Col-0) were sown on Murashige and Skoog medium with 0.5% gellan gum. After incubation for 2 days at 4°C in dark, the seeds were cultivated at 22°C on a 12:12-h light/dark cycle. The whole aerial part of plants 7 or 22 days after germination was collected 1, 7, 13, and 19 h after the start of light period and immediately frozen in liquid nitrogen and stored on −20°C until RNA extraction. Each individual plant was used as a sample for RNA-seq. Total RNA was extracted with the Maxwell 16 LEV Plant RNA kit (Promega, Madison, WI, USA). RNA-seq library preparation was performed as previously described (Nagano *et al.*, 2015); seven lanes of single-end 50-bp sequencing of the library were analyzed using the Hiseq2000 and HiSeq2500 systems (Illumina, San Diego, CA, USA). Sequences were pre-processed, mapped, and quantified according to a previously described pipeline (Kamitani *et al.*, 2016). Fastq files were deposited into the DNA Data Bank of Japan Sequence Read Archive as accession no. DRA005887.

### 4.2 Analysis of gene expression level variation bias

Owing to technical limitations, there was a time lag of several to 10 min during the harvesting of *Arabidopsis* leaf samples, potentially introducing a bias in the expression levels of some genes with respect to harvest time. In order to evaluate the variation bias in gene expression levels under each condition, we calculated the average gene expression levels from half of the samples harvested at early time points and half harvested at late time points. Gene expression level was regarded as stationary (unbiased) if the P value in the t test was > 0.2.

### 4.3 Cluster analysis

k-Means cluster analysis using R software (<u>http://www.r-project.org</u>) was performed for normalized RED profiles. The number of clusters was selected so as to minimize the Bayesian information criterion.

### 4.4 Data sources for gene classification

To classify each gene, the Gene Ontology Slim classification list was obtained from TAIR (<u>http://www.arabidopsis.org</u>). Data on essential genes were obtained from the SeedGenes Project (<u>http://www.seedgenes.org/GeneList</u>) (Meinke et al., 2008).

## Acknowledgments

The authors thank F. Kobayashi for assistance in RNA preparation, and H. Kudoh and S. Takada for helpful discussions. This research was partly supported by the Platform Project for Support in Japan Agency for Medical Research and Development (to A.A.); a Grant-in-Aid for Scientific Research on Innovative Areas “Integrated Analysis of Strategies for Plant Survival and Growth in Response to Global Environmental Changes” from the MEXT of Japan (no. 25119718 to A.A.); a Grant-in-Aid for Scientific Research on Innovative Areas “Initiative for High-Dimensional Data-Driven Science through Deepening of Sparse Modeling” from the MEXT of Japan (no. 26120525 to A.A.); MEXT KAKENHI Grant Number 17K05614 (to A.A.); and Japan Science and Technology Agency Core Research for Evolutional Science and Technology (no. JPMJCR15O2 to A. J. N.); and Japan Society for the Promotion of Science KAKENHI (nos. JP16H06171 and JP16H01473 to A. J. N.).

